# Pinpointing the neural signatures of single-exposure visual recognition memory

**DOI:** 10.1101/2020.07.01.182881

**Authors:** Vahid Mehrpour, Travis Meyer, Eero P. Simoncelli, Nicole C. Rust

**Affiliations:** Department of Psychology, University of Pennsylvania; Howard Hughes Medical Institute and Center for Neural Science, New York University

## Abstract

Memories of the images that we have seen are thought to be reflected in the reduction of neural responses in high-level visual areas such as inferotemporal (IT) cortex, a phenomenon known as repetition suppression (RS). We challenged this hypothesis with a task that required rhesus monkeys to report whether images were novel or repeated while ignoring variations in contrast, a stimulus attribute that is also known to modulate the overall IT response. The monkeys’ behavior was largely contrast-invariant, contrary to the predictions of an RS-inspired decoder, which could not distinguish responses to images that are repeated from those of lower contrast. However, the monkeys’ behavioral patterns were well-predicted by a linearly decodable variant in which the total spike count was corrected for contrast modulation. These results suggest that the IT neural activity pattern that best aligns with single-exposure visual recognition memory behavior is not RS but rather “sensory referenced suppression (SRS)”: reductions in IT population response magnitude, corrected for sensory modulation.

**Significance statement:** Memories of whether an image has been seen before are reflected in high-level visual cortex as “sensory referenced suppression (SRS)”: reductions in population response magnitude, corrected for sensory modulation.

## Introduction

Under the right conditions, we are very good at remembering the images that we have seen: we can remember thousands of images after viewing each only once and only for a few seconds ^1, 2^. How our brains support this remarkable ability, often called ‘visual recognition memory’ ^3^, is not well understood. The most prominent proposal to date suggests that memories about whether images have been encountered before are signaled in high-level visual brain areas such as inferotemporal cortex (IT) and perirhinal cortex via adaptation-like reductions of the population response to repeated as compared to novel stimuli, a phenomenon referred to as *repetition suppression* (RS) ^4–9^. Repetition suppression exhibits the primary attributes needed to account for the vast capacity of single-exposure visual memory behavior: response decrements in subsequent exposures are selective for image identity (even after viewing an extensive sequence of other images), and last for several minutes to hours ^5, 6, 10^. RS has also been shown to account for behavior in an image recognition memory task: a linear decoder with positive weights can predict single-exposure visual recognition memory behavior from neural responses in IT cortex ^10^.

Despite the fact that the RS hypothesis is consistent with available evidence, it seems likely to be too simplistic an explanation for visual recognition memory encoding. In particular, it is well-known that sensory neurons such as those of IT cortex are modulated not only by image memory, but also by stimulus properties such as image contrast ^11^. It is thus unclear whether and how these stimulus-induced effects interfere with judgments of whether images are novel or have been encountered before, and if they do not, how image memory can be decoded from neural responses in a way that disambiguates it from changes in these stimulus properties. To investigate this, we measured behavioral and neural responses of monkeys trained to report whether images were novel or repeated while disregarding image contrast (Fig. 1a).

**Fig. 1.**
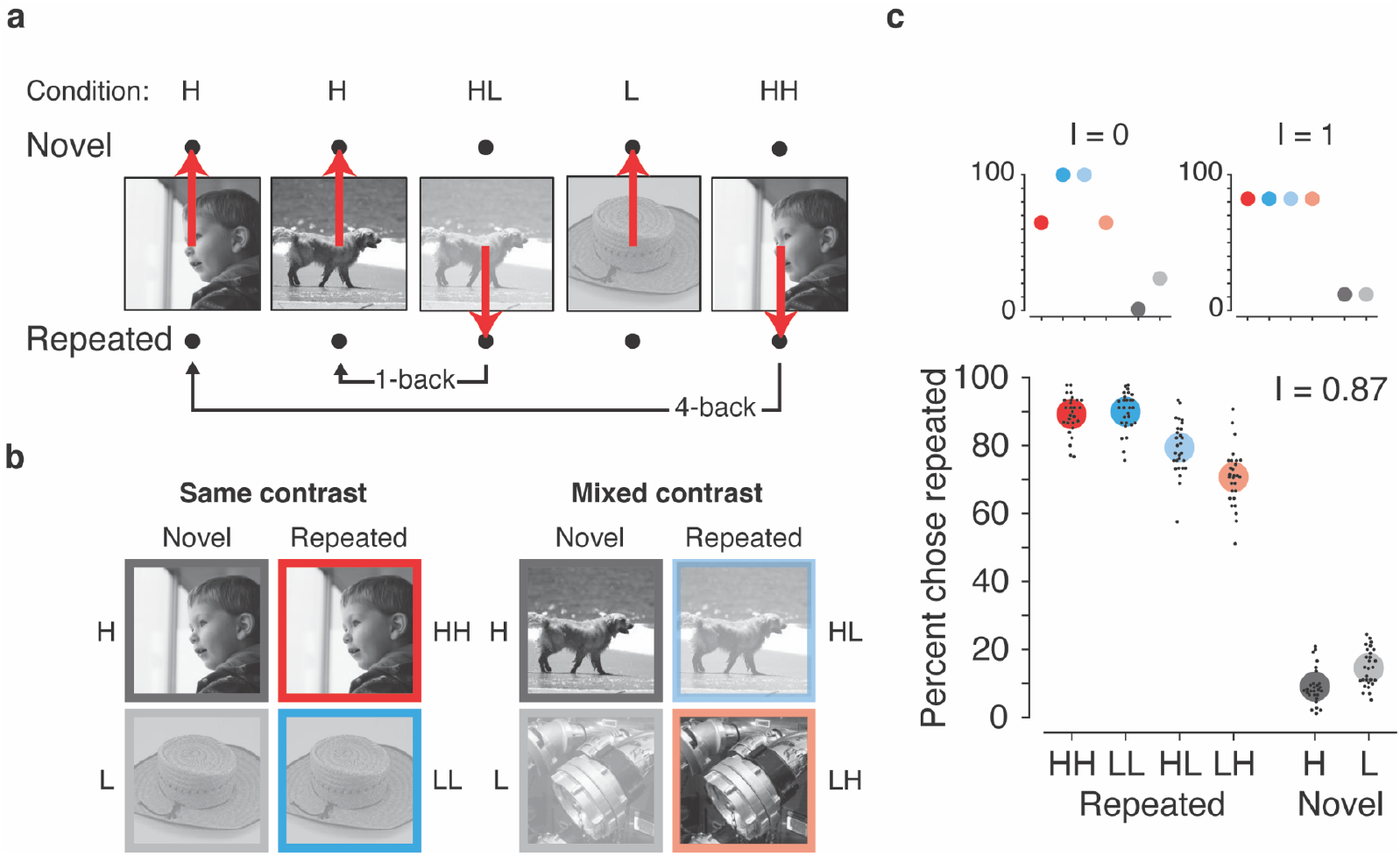
Visual memory behavior. **(a)** The contrast-invariant, single-exposure visual memory task. The monkeys viewed a sequence of images and reported whether they were novel (never seen before) or repeated (seen exactly once) while ignoring randomized changes in contrast. Monkeys were trained to saccade to one of two response targets to indicate their choice (red arrows). Images were repeated with a randomly chosen delay between the first and repeated presentation (‘n-back’). **(b)** Images were displayed at one of two contrast levels, yielding two conditions for novel images, high (H) and low (L), and four conditions for repeated images: HH (repeated H preceded by novel H), LL (repeated L preceded by novel L), HL (repeated L preceded by novel H), LH (repeated H preceded by novel L). The four repeated conditions were organized into “same-contrast” and “mixed-contrast” groups depending on whether the initial and repeated presentations were at the same or different contrasts, respectively. **(c)** Behavioral performance for the data pooled across monkeys in the task, where small black dots indicate average performance for an individual session and large colored dots indicate the average performance across sessions. A measure of contrast invariance, I, was computed as the ratio of the variance across contrast conditions and the variance with respect to the maximally contrast modulated pattern after taking overall performance into account, subtracted from one (see Methods). Insets illustrate the expected behavioral pattern with minimal (I = 0) and maximal (I = 1) contrast invariance.

## Results

### The contrast-invariant visual memory task

Monkeys viewed sequences of grayscale images, each presented for 500 ms, and each presented exactly twice (initially novel, then repeated). Novel and repeated images were presented with equal probability in all possible combinations of high (H) and low (L) contrasts, including (novel, repeated): HH, LL, HL, LH. We refer to the former two cases as the “same-contrast” conditions and the latter two as the “mixed-contrast” conditions (Fig. 1b). Monkeys were trained to report, on each trial, whether the observed image was novel or repeated, while disregarding image contrast (Fig. 1a). Here we refer to the change in state between novel and repeated trials as ‘image memory’. After training, the monkeys were largely able to disambiguate image memory from changes in image contrast: they performed equally well for both same-contrast conditions, and they were only modestly impaired for the mixed-contrast conditions (Fig. 1c). We quantified the degree of contrast invariance in the behavioral patterns with a measure in the range 0-1, where 1 indicates a behavioral pattern that is perfectly contrast invariant and 0 corresponds to the pattern that is maximally contrast dependent after taking into account the monkeys’ overall performance in each memory condition (see Fig. 1c insets). Behavioral contrast invariance values were high (combined data: 0.87; monkey1: 0.95, monkey2: 0.84; *SI Appendix*, Fig. S1), indicating that the monkeys were able to judge image memory while largely disregarding image contrast.

### RS and optimally weighted linear decoders fail to predict behavior

As the monkeys performed the task, we recorded neural responses in IT. Because accurate estimates of population response magnitude require many hundreds of units, data were concatenated across sessions into a larger pseudopopulation in a manner that combined trials within the same experimental condition (see Methods). Spikes were counted in a window starting 100 ms after stimulus onset (to allow for the latency of visual signals arriving in IT) and ending 400 ms later, at the termination of the image viewing period. The resulting pseudopopulation contained the responses of 856 units to 180 images each presented twice, and distributed evenly (and randomly) within the four conditions (i.e. 45 images for each of HH, LH, HL, LL). As an initial, summary analysis of the IT neural data, we quantified the magnitudes of both memory and contrast modulations as proportional reductions in the overall grand mean firing rate from the novel H condition to the repeated HH and novel L conditions for memory and contrast, respectively. Modulations for memory and contrast were 6% versus 3% when applied to the raw responses, and 13% versus 7% after subtracting out the pre-stimulus onset baseline.

Next, we assessed the hypothesis that RS of IT responses can explain visual memory behavior. We instantiated this hypothesis with a total spike count decoder, in which image memory was determined by comparing the total spike count with a threshold. We quantified the degree of alignment between neural predictions of behavioral patterns and the monkeys’ actual behavior with a measure termed ‘prediction quality (PQ)’. PQ was quantified as the normalized mean squared error between the actual behavioral patterns and best-fitting neural prediction of behavior (see Methods). The upper bound for our measure, PQ = 1, reflects a neural prediction that perfectly replicates the actual behavioral pattern, and PQ = 0 reflects the worst possible predicted behavioral pattern that was matched in overall performance (e.g., a pattern that was modulated entirely by changes in contrast, analogous to the insets in Fig. 1c). This ‘RS’ decoder produced a behavioral prediction that reflected confusions between changes in image memory with changes in contrast, for both repeated as well as novel images (compare with the insets in Fig. 1c) and low PQ (PQ_RS_ = 0.40; Fig. 2a). A control analysis confirmed that the predicted behavioral patterns on repeated trials were not consequence of misclassifications of those images when they were presented as novel (*SI Appendix*, Fig. S2), consistent with the interpretation that these behavioral patterns reflect confusion with contrast as opposed to other factors.

**Fig. 2.**
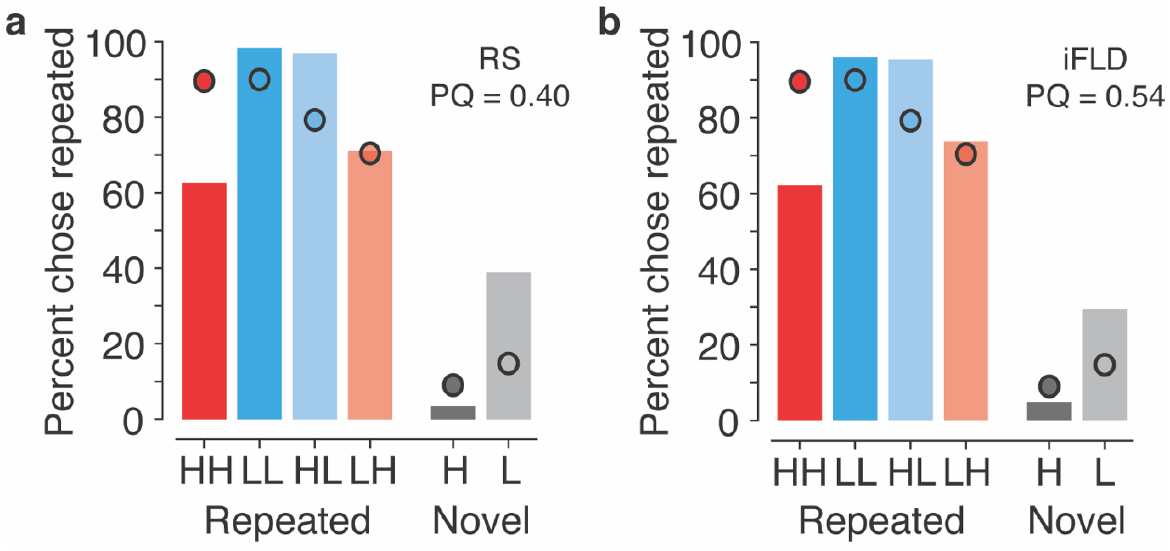
Traditional linear decoders confuse image memory and contrast and fail to map IT neural responses to behavior. Each panel reflects the monkeys’ actual behavioral patterns (dots) along with the predictions of a linear decoder applied to the recorded neural population (bars). (**a**) Total spike count decoder, motivated by RS. **(b)** Optimally weighted linear decoder, iFLD. Prediction quality (PQ) quantifies similarity between the neural predictions of behavior and the monkeys’ actual behavioral patterns (see Text).

The RS decoder is a linear decoder with uniform weighting over the neural population, so we wondered whether more carefully chosen weights might yield a linear decoder that could match the behavioral responses. Specifically, an optimally-weighted linear decoder was previously shown to be effective at aligning IT neural responses with visual memory behavior in the absence of contrast modulation ^10^. We used this same Fisher Linear Discriminant, computed assuming independence of neural responses, that weights each unit proportional to its memory discriminability, d’ (iFLD; see Methods). The iFLD differs from RS in that it weights each unit according to the amount of task-relevant information that it carries, and these weightings are signed: any units that exhibit repetition enhancement (on average) would be appropriately combined (with opposite sign) with units that exhibit repetition suppression. Despite the fact that this decoder is optimized to extract image memory information while disregarding contrast, we found that the iFLD also confused changes in image memory with changes in image contrast, and behavioral predictions were only slightly improved relative to RS (PQ_iFLD_ = 0.54; Fig. 2b). Poor behavioral predictions for RS and iFLD were replicated for each monkey individually (*SI Appendix*, Fig. S3; monkey 1: PQ_RS_ = 0.61, PQ_iFLD_ = 0.66; monkey 2: PQ_RS_ = 0.19, and PQ_iFLD_ = 0.53). We return to examine the underlying reasons for this failure below, in Fig. 5.

### Sensory referenced suppression is a good predictor of behavior

We wondered whether the monkeys’ behavioral patterns could be explained by an alternative linear decoder applied to the IT population responses. Given the substantial evidence in support of the repetition suppression hypothesis, we reasoned that the brain might be acting on a variant of this neural signature in which it corrects for the ambiguities in total spike count that are introduced by changes in contrast. Because this hypothetical decoding scheme operates by estimating and correcting for modulations in the total spike count due to variations in memory-irrelevant sensory attributes, we refer to this hypothesis as “sensory referenced suppression (SRS)”.

What would be required for SRS to be an effective account of the mapping of IT neural signals to behavior, if such a decoding scheme were restricted to act only on the IT population response? The fact that the total spike count is affected by contrast implies that the optimal linear decoder for contrast must at least partially overlap with (i.e. be nonorthogonal to) the RS decoding axis (a vector of ones, representing equal weights for each unit). We found that this was indeed the case: when applied to the pooled data, an optimized decoding vector for contrast lies in a direction 69° from the total spike count vector (labeled RS), indicating that information about contrast was largely non-overlapping but not orthogonal to RS (Fig. 3a). Next, we considered the family of linear discrimination vectors that live on the two-dimensional plane defined by RS and the contrast decoder. On this plane, we defined the direction of the RS decoder (with no contrast correction) as 0° (see Fig. 3a, top inset: ‘RS’). A decoding vector that is rotated clockwise from the RS decoder (away from the contrast decoder) in this plane is less affected by stimulus contrast. Rotation toward the contrast decoder exacerbates contrast modulation in the predicted behavioral patterns. Within this family of linear decoding schemes, we define SRS as the decoder that is orthogonal to (i.e. 90° from) the contrast decoder, and consequently minimizes contrast modulation in the neural prediction of behavioral patterns. The SRS was −21° from RS for the data pooled across both monkeys (Fig. 3b) and −23° and −18° for individual animals (*SI Appendix*, Fig. S3).

**Fig. 3.**
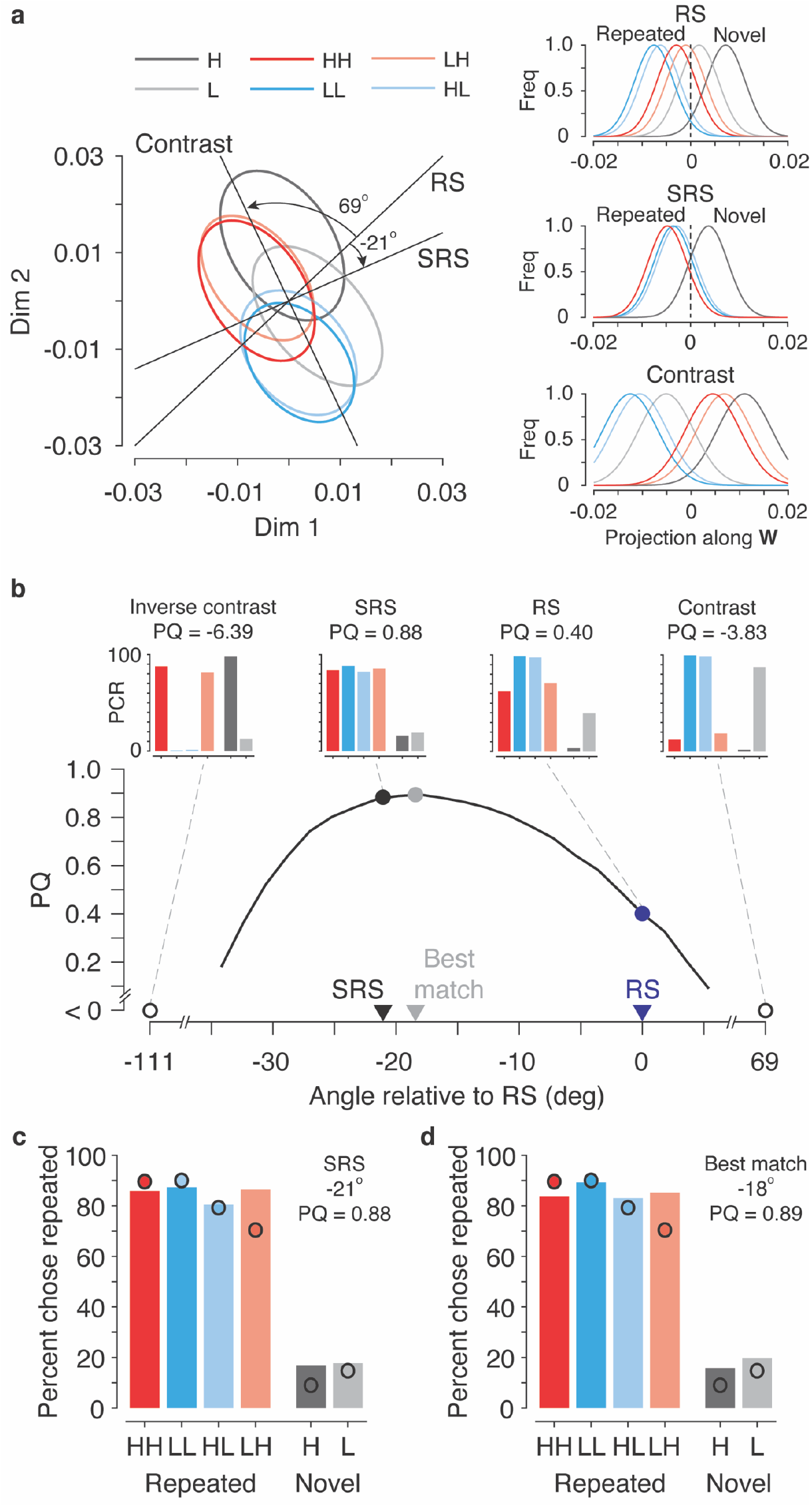
Neural predictions of behavior for a family of weighted linear decoders that include RS, an optimized contrast decoder, and SRS. (**a**) Projections of IT neural response distributions for all 6 stimulus conditions onto the 2-D plane defined by weight vectors for the total spike count vector (‘RS’, which uses a weight vector of all ones) and for a linear decoder optimized for contrast (‘Contrast’). Ellipses depict 95% probability intervals for 2-D histograms of the projection of neural responses onto this plane (see Methods). Insets show 1-D histograms of the projections of the distributions onto the three linear decoders. (**b**) The quality of the neural predictions of monkeys’ behavioral patterns (PQ) for the family of linear decoders that lie within the plane. Negative PQ values reflect predicted behavioral patterns that could not be rescaled to match overall performance because one or more entries were pinned at saturation (e.g., as a consequence of extreme contrast modulation). Each decoder corresponds to a rotation of the total spike count decoder, or equivalently, the weighted combination of the total spike count decoder and the contrast decoder. Markers indicate: SRS (black), which has minimal contrast sensitivity (i.e., orthogonal to the contrast axis); RS (blue), the total spike count decoder with no contrast correction; and the best behavioral match (maximal PQ – gray). Insets above depict the corresponding neural predictions of behavior. (**c-d**) The alignment of the monkeys’ actual behavioral patterns (dots) and the neural predictions of behavior (bars) for (**c**) SRS and (**d**) the decoder with the best behavioral match.

The SRS linear decoder provides good predictions of the monkeys’ behavioral patterns, both for the pooled data (PQ_SRS_ = 0.88; Fig. 3c), and for each monkey individually (monkey 1: PQ_SRS_ = 0.87; monkey 2: PQ_SRS_ = 0.93; *SI Appendix*, Fig. S4). It also provided a much better prediction of behavior than RS or the iFLD (pooled data: PQ_RS_ = 0.40 & PQ_iFLD_ = 0.54; monkey 1: PQ_RS_ = 0.61 & PQ_iFLD_ = 0.66; monkey 2: PQ_RS_ = 0.19 & PQ_iFLD_ = 0.53). These results suggest that SRS provides a considerably better description of the relationship between IT neural activity and behavior than RS or iFLD under the challenge of sensory-induced variations in population response magnitude (i.e. contrast modulation).

The SRS decoder can be thought of as isolating image memory information by correcting the IT population response for contrast modulation. We also considered a variant decoding scheme in which the contrast correction was applied by estimating and then subtracting contrast modulation from the RS decoder response (see Methods). The behavioral pattern predicted by this decoder reflected a higher degree of contrast modulation, and was less well aligned with behavior (PQ = 0.75; *SI Appendix*, Fig. S6) than SRS (PQ_SRS_ = 0.88; Fig. 3c).

The results presented thus far were obtained using response values averaged over a temporal window from 100-500 ms following stimulus onset. We found similar results when the analysis was applied to shorter time windows placed at different positions relative to stimulus onset. These analyses revealed that contrast modulation preceded memory modulations in IT, and was reflected throughout the entire viewing period following the initial latency delay (*SI Appendix*, Fig. S5a-b). Consequently, PQ_SRS_ rose and then remained high until the end of the viewing period and PQ_SRS_ was higher than PQ_RS_ for all time windows (*SI Appendix*, Fig. S5b).

### The SRS decoder had better image memory performance than RS

To better understand how memory and contrast were reflected in IT during these experiments, we shifted our focus away from the alignment between decoding predictions and behavior and toward overall performance for decoding image memory. These issues are best conceptualized by considering discriminability, rather than percent correct, as a measure of performance. Discriminability (often labelled d’ in the perceptual literature) is defined as the ratio of the difference between the means of the novel and repeated distributions divided by the square root of the average variance of those distributions (Fig. 4a). In our experiments, the variance of each distribution can be further decomposed into two components: (1) modulations within each distribution by contrast (Fig. 4a, ‘Contrast Var’), and (2) combined modulations arising from image identity and trial-to-trial variability (which cannot be dissociated, due to the single-trial nature of these experiments; Fig. 4a, ‘ID Var’) - see Methods.

**Fig. 4.**
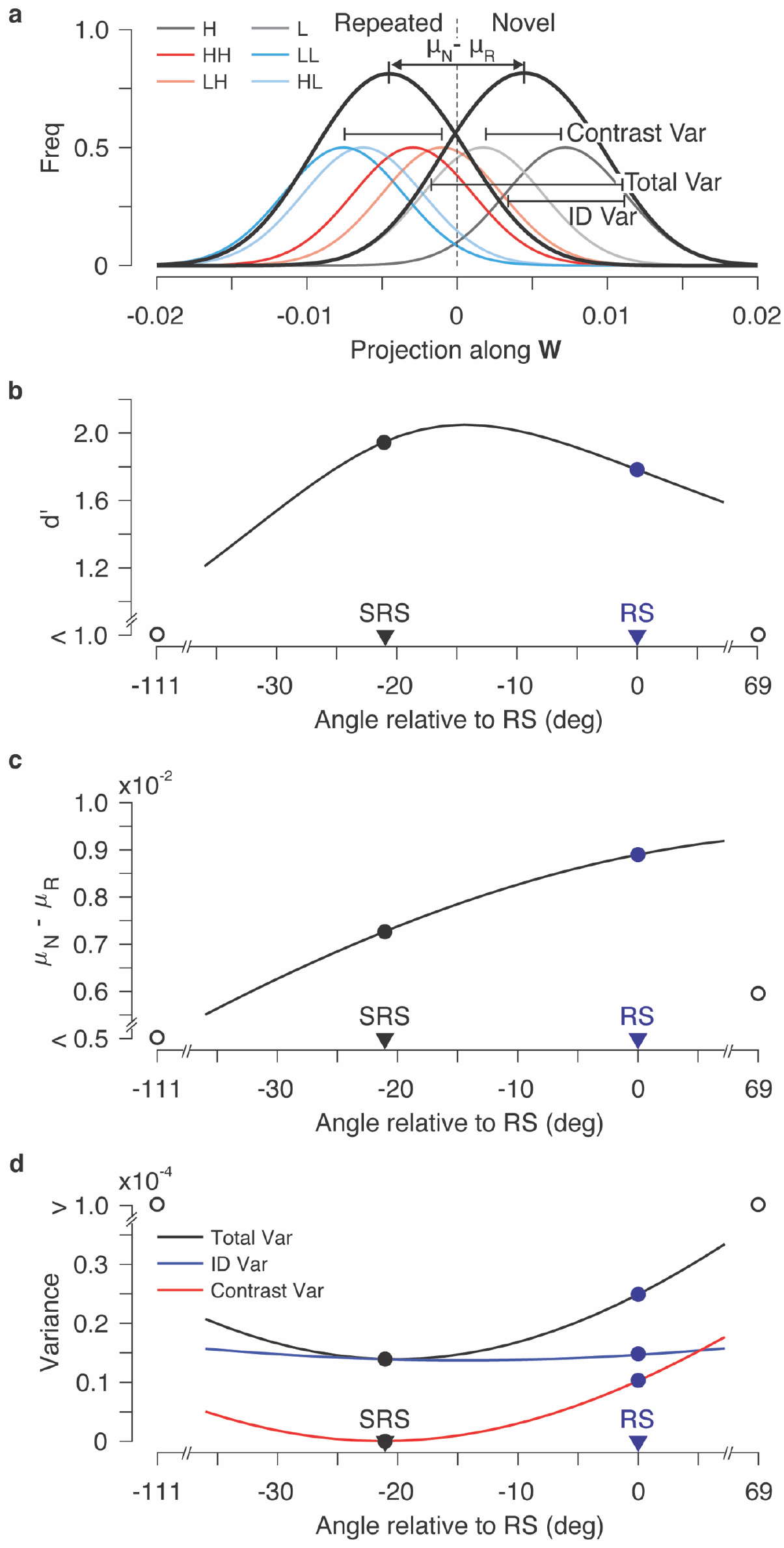
The population geometry impacting overall image memory performance for SRS and RS. (**a**) A schematic of linear decoder performance, computed as d’, for this task. Shown are 1-D histograms of the projection of the IT population responses onto a linear decoding axis **W.** Discriminability for image memory (d’) is computed as the difference between the means of novel and repeated distributions (*μ*_*N*_ − *μ*_*R*_) divided by the square root of the average total variance (total Var). (**b**) d’ as a function of angle on the 2-D plane defined in Fig. 3a. (**c-d**) Decomposition of d’: **(c)** the numerator (difference between means), and (**d**) the square of the denominator, the total variance (Total Var), further broken down into the variance due to image identity and trial-to-trial variability (ID Var) and contrast modulation (Contrast Var) – see Methods. In b-d, open circles at the right side of each graph indicate the values for projections along the contrast decoder.

**Fig. 5.**
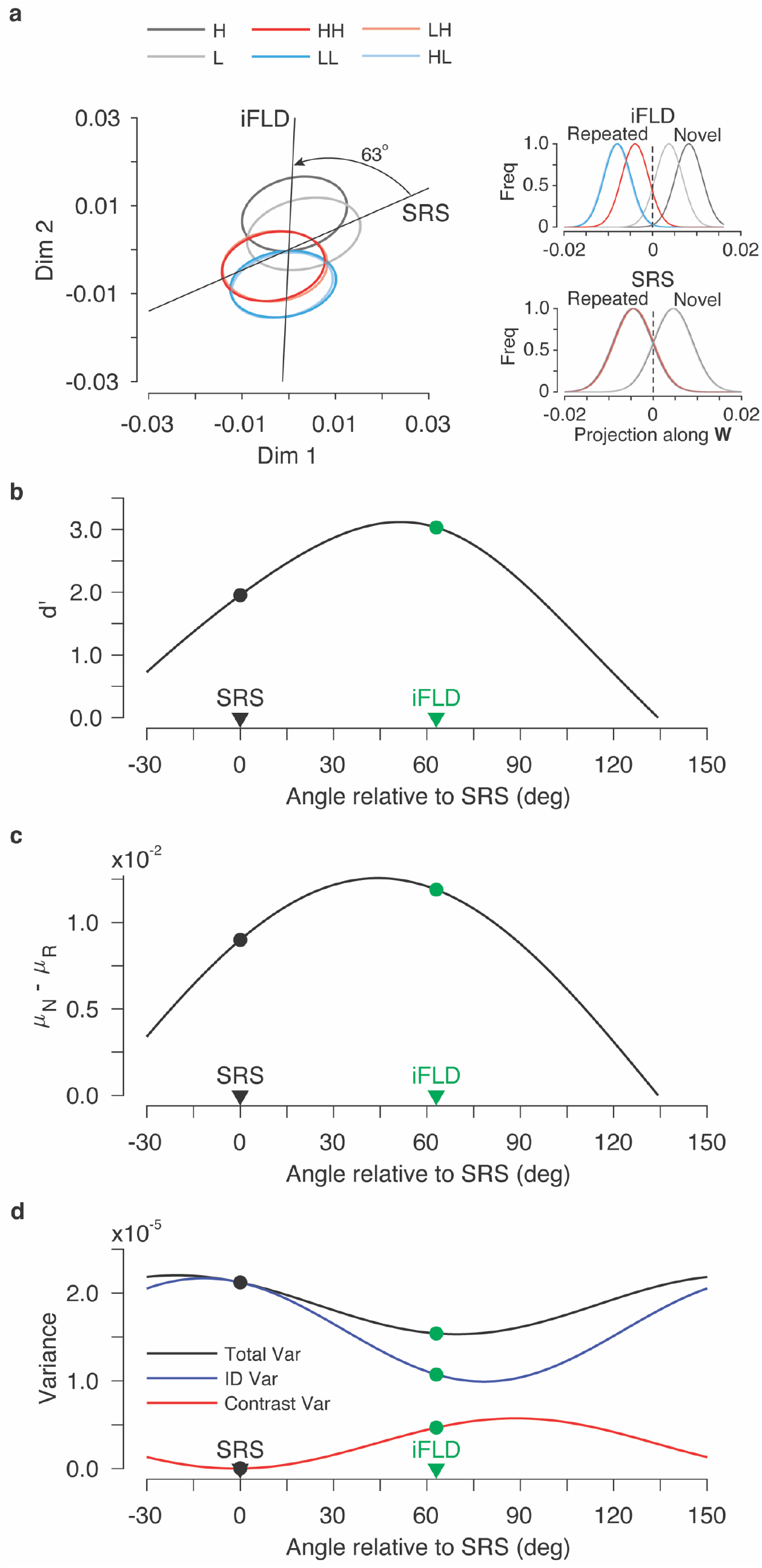
The population geometry impacting overall performance for SRS and iFLD. To explore population geometry absent the constraints imposed by limited samples, a model was fit to each unit and model parameters were used to create synthetic data. **(a)** Projections of the synthetic data onto the 2-D plane defined by SRS and a linear decoder optimized for memory, ‘iFLD’. Ellipses depict 95% probability intervals for 2-D histograms of the projection of neural responses onto this plane. Insets show 1-D histograms of the projections onto each linear axis. **(b)** d’ as a function of angle relative to SRS on the 2-D plane defined in panel a. **(c-d)** Decomposition of d’ into **(c)** It’s numerator, the difference between the means of the novel and repeated distributions and **(d)** The square of its denominator, the total variance (Total Var), further broken down into the variance due to image identity and trial-to-trial variability (ID Var) and contrast modulation (Contrast Var). In b-d, values corresponding to SRS and iFLD are labeled by black and green markers, respectively.

We found that, in addition to being a better predictor of behavior (Fig. 3b), the SRS decoder also had higher image memory performance than RS (Fig. 4b). This occurred despite the fact that novel and repeated means were actually closer together in the SRS direction than the RS direction (Fig. 4c). These decreases in mean separation were offset by decreases in variance (denominator of d’), plotted in Fig. 4d. These decreases in variance could in turn be attributed entirely to the elimination of contrast modulation. In sum, the superior performance of SRS resulted from novel and repeated distributions whose means were slightly closer together, but whose variances decreased even more as a consequence of eliminating contrast modulation along the SRS linear decoding axis.

### Relationship between SRS and iFLD decoders

The results presented above demonstrate that while the largely contrast-invariant patterns reflected in monkeys’ behavior are consistent with the SRS decoder (Fig. 3c), a linear decoder optimized for image memory on our data (the iFLD) confuses image memory with changes in image contrast (Fig. 2b). What do these differences imply about the geometry by which image memory and contrast are reflected in IT? To address these questions, we turned to simulations, where issues about population geometry can be investigated absent the constraints imposed by finite samples. To perform these simulations, we began by fitting a model to each single unit that we recorded. For each IT unit, the distribution of the visually-evoked firing rate response over stimuli was modeled by an exponential function ^12^, image memory and contrast were modeled as multiplicative modulations of the visually-evoked response, and the trial-to-trial distribution of spike counts was modeled as an independent Poisson process (see Methods). The four parameters fit for each unit included: (1) mean firing rate (the mean of the exponential), (2) the visually-evoked tuning bandwidth, (3) image memory sensitivity, and (4) contrast sensitivity (see Methods). We found that ‘synthetic’ data from the resulting model population recapitulated all aspects of the physiological data that we have highlighted thus far, including contrast modulation in the RS predictions (*SI Appendix*, Fig. S7a, top inset), contrast-invariant SRS (*SI Appendix*, Fig. S7a, middle inset), and overall d’ that was higher for SRS than RS as a consequence of eliminating contrast modulation (*SI Appendix*, Fig. S7b-d).

Next, to understand the relationship between SRS and the iFLD, and why the iFLD did not exhibit contrast invariance, we performed a set of analysis similar to those described for Figs. 3–4 but within the plane spanned by SRS and iFLD (Fig. 5a). The iFLD is optimal (under the assumption of Gaussian-distributed independent response), and indeed has higher discrimination performance than SRS (Fig. 5b). Increased d’ for iFLD over SRS resulted from both an increase in the distance between the means of distributions for novel and repeated images (i.e. the d’ numerator; Fig. 5c) as well as a decrease in the variance of distributions for novel and repeated images (i.e. the d’ denominator; Fig. 5d). Intriguingly, the overall reduction in total variance along the iFLD axis relative to SRS resulted from an *increase* in contrast modulation that were offset by a larger decrease in identity modulation relative to SRS (Fig. 5d). This was because identity modulation and contrast modulation were anti-correlated on this plane: decreases in one (e.g. identity modulation) were accompanied by increases in the other (e.g. contrast modulation; Fig. 5d). In other words, the iFLD failed to predict contrast invariance in behavioral patterns because it could achieve higher image memory performance by reducing identity variance, which was anti-correlated with contrast.

To complement the 2-D plots presented in Fig. 4 (RS and SRS) and Fig. 5 (SRS and iFLD), *SI Appendix*, Fig. S8 depicts memory decoding performance in the 3-D space defined by SRS, RS and iFLD, for both the real and synthetic data.

## Discussion

Humans and nonhuman primates have a remarkable ability to remember the images that they have seen ^1, 2, 5, 6, 13^. It has been suggested that image memory is signaled in high-level visual brain areas such as IT via changes in population response magnitude, or repetition suppression (RS) ^4–9^. We have challenged this explanation, by examining neural and behavioral memory responses while independently manipulating image contrast. IT population response was modulated by contrast, but monkeys’ behavioral reports of image memory were largely invariant to changes in image contrast (Fig. 1), inconsistent with the RS hypothesis (Fig. 2a). Behavioral invariance also could not be reconciled with our previous work, which proposed that image memory is linearly decoded from IT by weighting each neuron proportional to the amount and sign of memory-relevant information that it carries ^10^ (Fig. 2b). However, the monkeys’ behavioral patterns were linearly decodable from IT (Fig. 3c), using a linear decoder that corrects the total spike count decoder by eliminating its contrast dependence. We refer to this linear decoding scheme as sensory referenced suppression ‘SRS’, because it can be understood as estimating image memory from the total spike count after correcting for sensory modulation (Fig. 3a).

The hypothesis that image memory is encoded in high-level visual cortex as RS has a mixed history, with some studies finding support for this hypothesis ^6, 8, 10, 14, 15^ and others finding evidence against it ^16, 17^. In an earlier study, we reported that some instantiations of RS decoders were good predictors of the rates of behavioral remembering and forgetting when tested with randomly selected images, under the assumption that all novel images evoke the same magnitude population response from IT ^10^. The work we present here suggests that modifications of RS are required to account for single-exposure visual memory behavior when factors other than image memory modulate the magnitude of the population response. That is, we find that when novel images evoke lower firing rates due to changes in contrast (e.g. L vs H), an RS classifier confuses those differences in firing rate due to contrast for differences in image memory, and predicts higher memory performance when those images are repeated (e.g. LL versus HH). These predicted patterns are at odds with the actual behavioral patterns of the monkeys, who exhibit similar performance for the LL and HH conditions. SRS resolves this discrepancy by removing the contrast dependency from the RS memory encoding scheme.

A number of factors other than contrast are known to modulate the IT population response, including stimulus attributes such as object size ^11^, a diverse set of stimulus attributes that contribute to image memorability ^18–20^, and external factors such as surprise ^21, 22^ and attention ^8, 23^. The SRS decoding scheme that we have proposed could, in principle, provide a mechanism for the brain to disambiguate image memory-induced changes in IT population response magnitude from changes due to the combination of all of these other factors. In principle, all that is required to generalize the SRS decoding scheme is for the effect of these other factors at least partially nonoverlapping with that of image memory. This would be expected when individual units have heterogeneous sensitivities for the multiple factors that modulate the population response. In that case, a linear decoder can be constructed by eliminating each of these factors in turn from the RS decoder, analogous to the successive orthogonalizations performed in the Gram–Schmidt procedure. Future work will be required to determine how such a decoding scheme might be learned by the brain.

What is the origin of the IT magnitude variation that aligns with single-exposure visual recognition memory behavior? RS is found at all stages of visual processing from the retina to IT, and it strengthens in both magnitude and duration as one ascends the visual cortical hierarchy ^24^. Consequently, a hierarchical cascade of feed-forward, adaptation-like mechanisms may underlie RS measured in IT ^25^. There are also indications that RS within IT may arise from changes in synaptic weights between recurrently connected units within IT itself ^25, 26^. Finally, a component of RS in IT is likely to be fed back to IT from higher brain areas such as perirhinal cortex or hippocampus. While the assertion that top-down processing contributes to RS in high-level visual cortex has been controversial ^25, 27–29^, recent evidence from a patient with medial temporal lobe (MTL) damage supports a role for feedback from MTL structures to RS in high-level visual cortex ^30^. Within one MTL structure, the hippocampus, single-exposure recognition memory behavior has been linked with RS ^31, 32^ as well as synchronizations between gamma oscillations and spikes^33^. However, because these evaluations were not made in a manner that challenges RS with other factors that affect response magnitude, additional work will be required to determine whether SRS is a better description than RS of the neural signatures that reflect single-exposure visual recognition memory behavior in MTL structures that lie downstream from IT. What is clear is that pinpointing the neural signatures that align with single-exposure visual memory behavior is a good first step toward understanding how the primate brain manages to remember images so remarkably well.

## Acknowledgments

This work was supported by the Simons Foundation (Simons Collaboration on the Global Brain award 543033 to NCR and 543047 to EPS), the National Eye Institute of the National Institutes of Health (award R01EY020851 to NCR), the National Science Foundation (CAREER award 1265480 to NCR), and the Howard Hughes Medical Institute (investigatorship to EPS).

## REFERENCES

### Citation Diversity Statement

Recent work in neuroscience and related fields has identified citation biases whereby work from women and minorities are under-cited relative to other papers in the field^34–36^. In crafting this manuscript, we sought to proactively consider citation bias. Following ref. ^34^, the gender balance of citations was quantified based on the first names of the first/last authors using open source code^37^. Excluding self-citations, this article contains 58.3% man/man, 16.7% man/woman, 19.4% woman/man, and 5.6% woman/woman citations. For comparison, proportions estimated from articles in the 5 top neuroscience journals (as reported in ref. ^34^) are 58.4% man/man, 9.4% man/woman, 25.5% woman/man, and 6.7% woman/woman.

## Methods

Experiments were performed on two adult male rhesus macaque monkeys (*Macaca mulatta)*, with weights of 11 kg and 12 kg and estimated ages of 10 years and 7 years, respectively. Both animals were implanted with head posts and recording chambers. All procedures were performed in accordance with the guidelines of the University of Pennsylvania Institutional Animal Care and Use Committee.

### The single-exposure, contrast-invariant visual memory task

All behavioral training and testing were performed using standard operant conditioning (juice reward), head stabilization, and high-accuracy, infrared video eye tracking. Stimuli were presented on an LCD monitor with an 85 Hz refresh rate using customized software (http://mworks-project.org).

As an overview of the monkeys’ task, each trial involved viewing one image for 500 ms, after which the monkeys indicated whether it was novel (never seen before) or repeated (seen exactly once prior) with an eye movement to one of two response targets. Images were never presented more than twice during the entire training and testing period of the experiment. Trials were initiated when the monkey fixated on a small red square (0.25°) on the center of a gray screen followed by a 200 ms delay before a 4° image appeared within a circular aperture, positioned at the center of gaze. The monkeys needed to maintain fixation of the stimulus for 500 ms, at which time the red square turned green (the go cue) and the targets appeared. The monkeys then made a saccade to a target indicating whether the stimulus was novel or repeated, and correct responses were rewarded with juice. Targets were positioned 8° above or below the stimulus. The association between the target (up vs. down) and the report (novel vs. repeated) was swapped between the two animals.

The images used in these experiments collected via an automated procedure that gathered images from the internet. Images smaller than 96*96 pixels were not considered. Eligible images were cropped to be square and resized to 256*256 pixels. Duplicate images were removed. Colored images were converted to grayscale and were presented at two contrasts (“low (L)” and “high (H)”) in all possible combinations as novel and repeated (novel-repeated = low-low (LL); high-high (HH); low-high (LH); high-low (HL)). Contrast modifications were applied in a manner that did not adjust image luminance (*L*_v_), the mean pixel intensity. Images with *L*_v_ outside the range 0.25 – 0.75 were excluded. The computation of contrast began by first computing the median of the pixel intensities that fell above and below *L*_v_, *L*_v-hi_ and *L*_*v*-lo_. The native contrast for each image C_native_ was computed as:

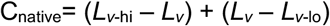

Each image was manipulated to produce a high contrast version (C_hi_ = 0.4) and low contrast version (C_lo_ = 0.2) via a procedure that maintained *L*_*v*_ for each image. Adjustments to contrast involved: 1) subtracting the mean pixel value, 2) rescaling the residual pixel values all by the same amount, and 3) adding back the mean. When the procedure resulted in the saturation of more than 10% of pixels beyond their maximal value (black and white), that image was excluded.

Trial locations for novel images and their repeats were presented with a uniform distribution of the subset of n-back used in the experiment. The n-back distribution was adjusted for each monkey based on training history to approximately equate overall performance between the two animals: n-back = 1, 4, 16, and 32 for monkey 1, and n-back = 1, 2, 4, and 8 for monkey 2. The specific random sequence of images presented during each session was generated offline before the start of the session. Uniform n-back distributions were achieved by constructing a sequence slightly longer than what was anticipated to be needed for the session, and by iteratively populating the sequence with novel images and their repeats at positions selected randomly from all the possibilities that remained unfilled. Because the longest n-back values (8 or 32) were the most difficult to fill, a fixed number of those were inserted first. In the relatively rare cases that the algorithm did not converge, it was restarted. The result was a partially populated sequence in which 83% of the trials were occupied. Next, the remaining 17% of trials were examined to determine whether they could be filled with novel/repeated pairs from the list of possible n-back options. The very small number of trials that remained after all possibilities had been extinguished (e.g. a 3-back scenario) were filled with ‘off n-back’ novel/repeated image pairs and these trials were disregarded in later analyses.

The monkeys’ behavioral patterns were computed for each condition after collapsing across n-back. The degree of contrast invariance reflected in each monkey’s session-averaged behavioral pattern was computed as the mean of contrast invariance computed for the novel and repeated memory conditions separately. Within each memory condition M, contrast invariance (I) of the behavioral pattern X in either memory condition was defined by:

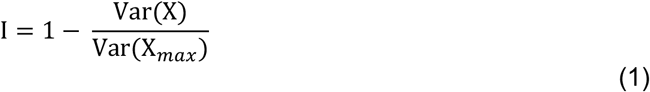

Where, Var(X) is the variance of pattern X, and Var(X_*max*_) is the maximum possible variance associated with contrast in memory condition M given monkeys’ overall performance in the same memory condition M. For example, the X_*max*_ for an overall performance across the repeated conditions of 85% would correspond to 70%, 100%, 100% and 70% for HH, LL, HL and LH, respectively.

### Neural recording

The activity of neurons in IT was recorded via a single recording chamber in each monkey. Chamber placement was guided by anatomical magnetic resonance images in both monkeys. The region of IT recorded was located on the ventral surface of the brain, over an area that spanned 5 mm lateral to the anterior middle temporal sulcus and 14-17 mm anterior to the ear canals. Recording sessions began after the monkeys were fully trained on the task and behavioral performance had plateaued. The depth and extent of IT was mapped within the recording chamber in a previous experiment ^1^. Combined recording and behavioral training sessions happened 2-5 times per week across a span of 4 weeks (monkey 1) and 6 weeks (monkey 2). Neural activity was recorded with 24-channel U-probes (Plexon, Inc.) with linearly arranged recording sites spaced with 100 μm intervals. Continuous, wideband neural signals were amplified, digitized at 40 kHz and stored using the Grapevine Data Acquisition System (Ripple, Inc.). Spike sorting was done manually offline (Plexon Offline Sorter). At least one candidate unit was identified on each recording channel, and 2-3 units were occasionally identified on the same channel. Spike sorting was performed blind to any experimental conditions to avoid bias. A multi-channel recording session was included in the analysis if: (1) the recording session was stable, quantified as the grand mean firing rate across channels changing less than 3-fold across the session; (2) over 50% of neurons were visually responsive (a loose criterion based on our previous experience in IT), assessed by a visual inspection of the rasters; and (3) the number of successfully completed novel/repeated pairs of trials exceeded 100. In monkey 1, 19 sessions were recorded and five were removed (one based on criterion 1 and four based on criterion 3). In monkey 2, 15 sessions were recorded and one was removed (based on criterion 1). The resulting data set included 14 sessions for monkey 1 (n = 427 candidate units), and 14 sessions for monkey 2 (n = 429 candidate units). The sample size (number of successful sessions recorded) was chosen based on our previous work ^1^.

The data reported here correspond to the subset of images for which the monkeys’ behavioral reports were recorded for both novel and repeated presentations (e.g. trials in which the monkeys did not prematurely break fixation during either the novel or the repeated presentation of an image). Accurate estimate of population response magnitude requires many hundreds of units, and when too few units are included, magnitude estimates are dominated by the stimulus selectivity of the sampled units. To perform our analyses, we thus concatenated units across sessions to create larger pseudopopulations. When creating these pseudopopulations, we aligned data across sessions in a manner that preserved whether the trials were presented as novel or repeated and their experimental contrast condition. To prevent artificial correlations from influencing our results, analyses were performed after re-randomizing the responses within each condition for each unit to create many pseudopopulations. To deal with varying data sizes across sessions, the number of images included in the analysis was selected to balance incorporating data of equal sizes across sessions with not needlessly discarding data. NaNs were used as place holders for the more limited sessions in which data did not exist. The resulting pseudopopulations consisted of the responses to 180 images presented as both novel and repeated (i.e. 45 images per condition: HH, LL, HL, and LH). Spikes were counted in a temporal window over the range 100-500 ms following stimulus onset.

#### Contrast and memory modulations

Contrast (*c*) and memory (*m*) modulations were computed from the grand mean firing rate (GMFR) across all units and images as:

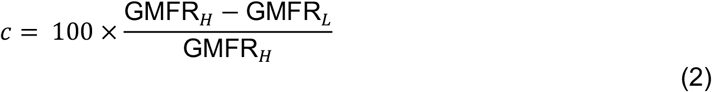

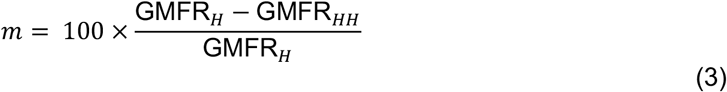

In the Results, we present both raw and baseline-corrected contrast and memory modulations. To determine the modulations after correcting for pre-stimulus baseline activity, the pre-stimulus GMFR in a 200-ms pre-stimulus time window was subtracted from the GMFR for each condition.

### Linear population decoders

For all decoders, the population response was quantified as the vector 6 containing spike counts on a given trial. To ensure that the decoder did not erroneously rely on visual selectivity, the decoder was trained on balanced pairs of novel/repeated trials in which monkeys viewed the same image (regardless of behavioral outcome or experimental contrast condition).

#### Cross-validated training and testing

We applied the same, iterative cross-validated procedure for each linear decoder. On each iteration of the resampling procedure, the responses for each unit were randomly shuffled within each experimental condition to ensure that artificial correlations (e.g. between the neurons recorded in different sessions) were removed. Each iteration also involved setting aside the responses to one randomly selected image within each contrast condition (presented as both novel and repeated, for 8 trials in total) for testing classifier performance. The remaining trials were used to train one of the linear decoders to distinguish novel versus repeated images invariant to contrast, where the novel and repeated classes included the data corresponding to all n-backs and all trial outcomes. A neural prediction of the proportion of trials on which “repeated” would be reported was computed as the proportion of each distribution that took on a value less than the criterion. Finally, the predicted response pattern was rescaled by a rescaling parameter (see below) as a proxy for adjusting the population size to consider.

All decoders in this study took the general form of linear discriminators. The class (novel/repeated) of a population response vector, 6 was determined by the sign of

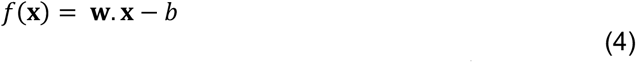

where **w** is an N-dimensional weight vector in the N-dimensional IT neural space (N is the number of units), and *b* is decision criterion, given by:

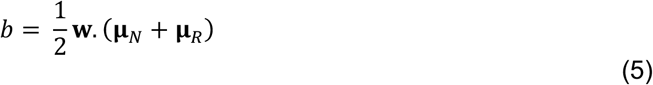

Here **μ**_*N*_ and **μ**_*R*_ are the mean population response vectors across novel and repeated images in the training set, respectively. A population response vector **x** was classified as “novel” if *f*(**x**) > 0, and “repeated” if *f*(**x**) < 0.

#### Spike count classifier (associated with repetition suppression, RS)

Arguably the simplest classifier, the total spike decoder uses a homogeneous weight vector:

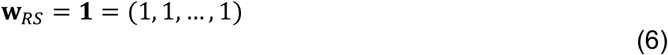

#### Fisher Linear Discriminant (iFLD)

The iFLD used in this study follows our previous implementation^1^. The Fisher Linear Discriminant (FLD) is defined as:

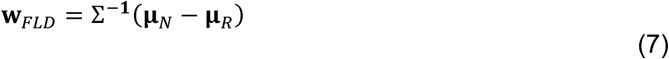

where Σ^−1^ is the inverse of the average covariance matrix across novel and repeated conditions:

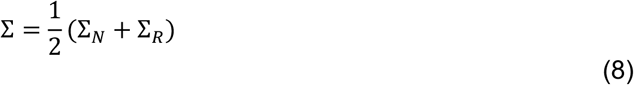

The dimensionality of our neural populations is high enough that we do not have enough data to obtain reliable covariance estimates (the amount of data needed for acceptable estimates of the off-diagonal entries is >10x what we collected in a single session). As such, we assume independence of the stimulus responses within conditions (i.e., we set the off-diagonal entries to zero). The resulting iFLD uses a weight for each unit that is proportional to its visual memory discriminability (d’):

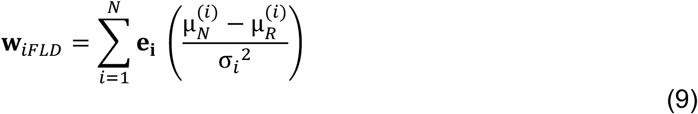

where **e**_**i**_ is the unit vector along *i*-th dimension (*i*-th unit’s response); N is the number of units; 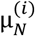 and 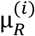 are *i*-th unit’s mean responses to novel and repeated images, respectively; and 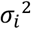 is the *i*-th unit’s average response variance across novel and repeated conditions:

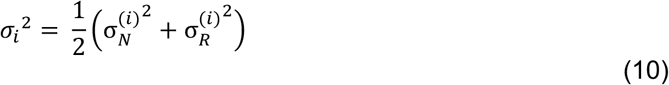

#### Family of contrast-corrected linear decoders

The family of contrast-corrected linear decoders are based on weight vectors that are rotated within the plane containing the RS decoder (**1**) and a contrast decoder, **W**_***c***_:

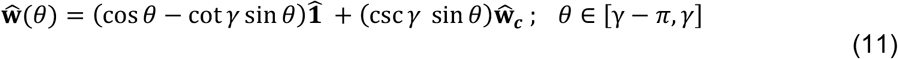

where, 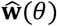, 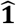 and 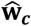 are the unit vectors representing the decoding axis, RS decoder, and contrast decoder, respectively. *θ* is the angle between the decoder axis 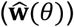 and the RS axis (**1**), and *γ* is the angle between the RS and contrast axes. The contrast weight vector **W**_***c***_ was defined as:

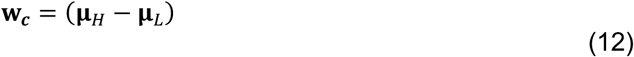

where **μ**_*H*_ and **μ**_*L*_ are the mean population response vectors across high and low contrast images in train set, respectively. This is a simple form of FLD that arises when the average covariance is a multiple of the identity, and is sometimes called a “prototype classifier”. We define the SRS decoder as the axis that is orthogonal to contrast, i.e.

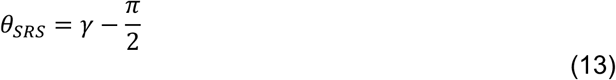

#### Variant decoding scheme that incorporates a contrast correction

We also evaluated a variant decoding scheme that corrected for contrast modulation (*SI Appendix*, Fig. S6). This decoder operated by correcting modulations caused by contrast along the RS axis by estimating and then subtracting the mean of population response across novel images for each contrast condition. The estimate of the mean population response at each contrast was computed after classifying novel images by contrast based on the training data, using the same prototype linear decoder used for SRS (Eq. 12).

#### Family of linear decoders in a 3D subspace spanned by SRS, RS, and iFLD

To compute the 3D plots presented in *SI Appendix*, Fig. S8, we considered a 3D subspace of our high dimensional neural space spanned by SRS, RS, and iFLD where plane **π**_1_: **w**_***SRS***_ ∧ **w**_***RS***_ intersects with plane **π**_2_: **w**_***SRS***_ ∧ **w**_***iFLD***_ along SRS (∧ denotes exterior product; see *SI Appendix*, Fig. S8a). In this 3D subspace a decoding axis is given by:

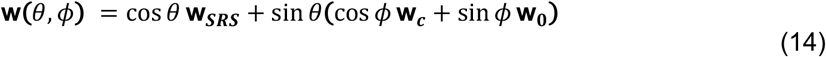

where *θ* and *ϕ* are polar and azimuthal angles in a spherical coordinate system measured relative to **w**_***SRS***_ (zenith direction) and **w**_***c***_ in the plane **π**_1_: **w**_***SRS***_ ∧ **w**_***RS***_ (azimuth reference), respectively. Furthermore:

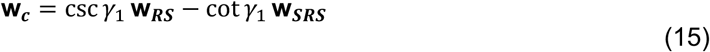

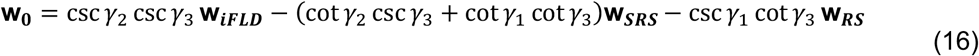

and

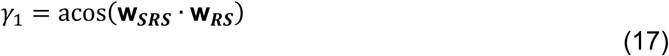

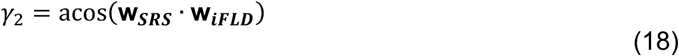

*γ*_3_ is the angle between planes **π**_1_: **w**_***SRS***_ ∧ **w**_***RS***_ and **π**_2_: **w**_***SRS***_ ∧ **w**_***iFLD***_. This angle can be determined in exterior algebra as:

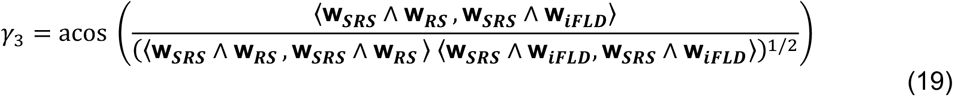

where

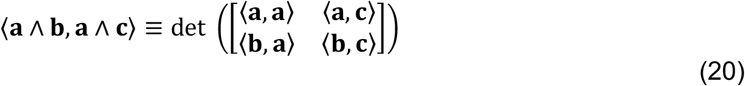

〈. , . 〉 and det(.) are the operators of inner product and determinant, respectively.

#### Rescaling parameter and prediction quality (PQ)

Comparing IT population decoding performance with behavior depends on the neural population size under consideration, and there is no good way to choose this *a priori.* We thus applied a fitting approach for each decoder. After confirming that performance using all recorded units in our dataset fell below saturation, we simulated increases in population size by fitting a single rescaling parameter (*α*) to minimize the MSE between the neural predictions and actual behavioral patterns. We emphasize that while this adjustment changed the overall performance, it did not impact the shape of the predicted behavioral patterns. The minimization of MSE yields the standard analytical solution for *α*:

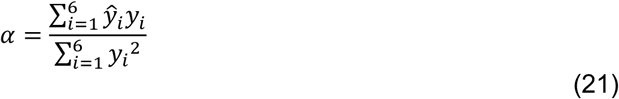

where 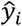 and *y*_*i*_ are the actual and neutrally predicted performance for condition i, respectively, and i indexes each of six conditions {HH, LL, HL, LH, H, L}.

Next, to quantify the quality of the fit after rescaling the predicted pattern, we computed a measure of prediction quality (PQ):

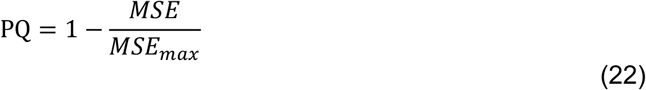

where *MSE* and *MSE*_*max*_ denote the mean squared error of the rescaled predicted pattern and the pattern with maximum MSE that was matched in overall performance, respectively, i.e.

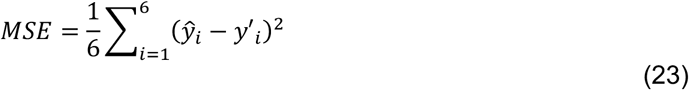

and

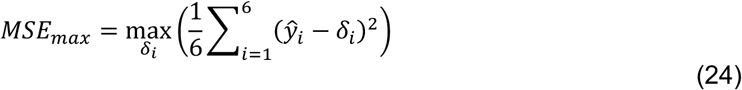

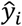 and *y*′_*i*_ are the actual and rescaled predicted performance for condition i, respectively, and i corresponds to each of six conditions including HH, LL, HL, LH, H, and L. Each *δ*_*i*_ was chosen to be either 1 or 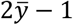 (with 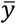 the mean performance across all six conditions), in order to maximize MSE. The upper bound of PQ = 1 reflects a neural prediction that perfectly replicates the actual behavioral pattern. A value of PQ = 0 reflects the worst possible predicted behavioral pattern that is matched in overall performance. Negative PQ values reflect predicted behavioral patterns that could not be rescaled with *α* to match overall performance because one or more entries were pinned at saturation (e.g., as a consequence of extreme contrast modulation).

#### Covariance error ellipse

Error ellipses (shown in Fig. 3a, 5a, and *SI Appendix*, Fig. S7a) were computed by first projecting the neural response vectors onto the non-orthogonal discriminant axes 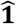 and 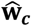, producing coordinates (u, v). These were transformed to orthogonal coordinates using a transformation matrix (derived from Eq. 11):

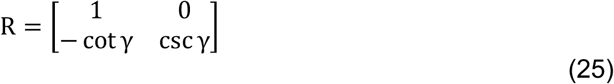

where γ is the angle between the two discriminant axes:

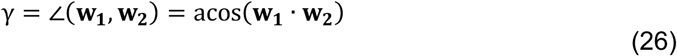

We then rotate this coordinate system in the plane by angle *φ*, using transformation matrix:

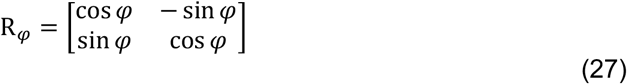

Combining Eq. 25 and 27 gives an expression for the (x, y) coordinates of the projected neural responses:

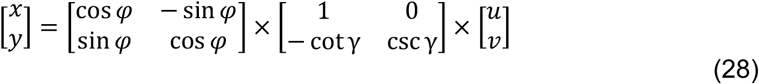

For each condition, the covariance matrix of the transformed data was computed, and the eigenvectors of this matrix provide the major and minor axes of the associated ellipse. To determine the dimensions of the ellipse, we multiplied the square root of the eigenvalues by a scale factor equal to the square root value of the cumulative chi-square distribution function (CDF) for 2-degrees of freedom evaluated at 95%.

#### Decomposition of total variance into variance due to identity/trial variability and contrast

For Figs. 4, 5, and *SI Appendix*, Fig. S7, we used the following equations to decompose the average variance across novel (N) and repeated (R) conditions, 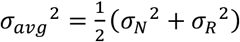, into the variance due to image identity and trial variability (ID) and contrast (C):

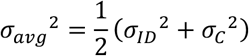

Where:

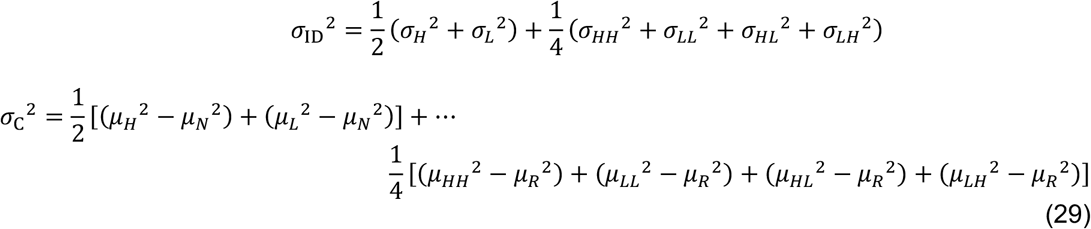

In each condition, *σ* and *μ* denote the standard deviation and mean of the corresponding distribution, respectively.

### Fitting the four-parameter tuning model to each unit and synthesizing data

In Fig. 5, and *SI Appendix*, Fig. S8 we assessed the population geometry in the limit of infinite samples by fitting a model to each unit that we recorded, and then using these models to synthesize population data. A 4-parameter model was used to describe the mean spike count response of each unit:

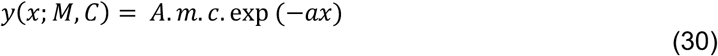

where x is stimulus rank, M is image memory condition (novel or repeated), C is image contrast (high or low), A is amplitude, m is memory modulation (set to 1 for novel images, and a fitted value for repeated images), c is contrast modulation (set to 1 for high contrast images, and fitted value for low contrast), and a controls stimulus selectivity.

We estimated each unit’s tuning curve parameters by maximizing the likelihood (MLE) of observing the spike count data from 100 ms to 500 ms relative to stimulus onset using the techniques introduced in ref. ^2^. If {*ν*_1_, *ν*_2_, … , *ν*_*n*_} is the spike count data for a unit in all six conditions (n trials in total), the log-likelihood of observing the data is given by:

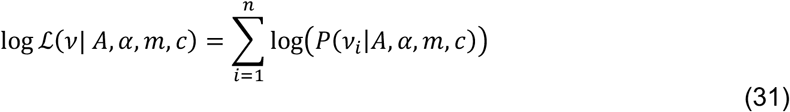

where

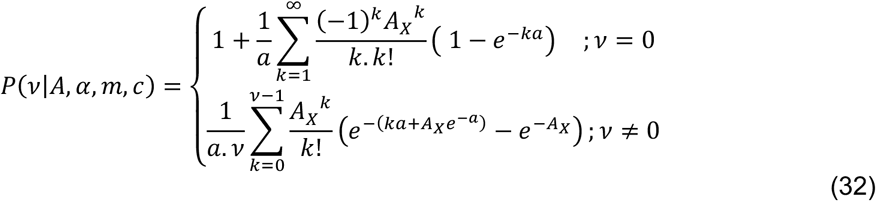

and

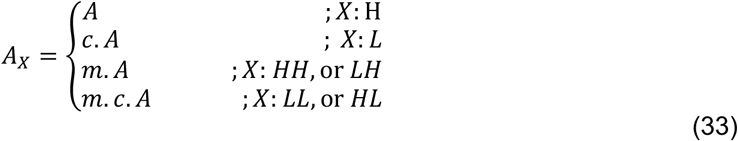

We estimated four tuning parameters of the unit by maximizing Eq. 31 with respect to the parameters A, a, m, and c.

Goodness of fit was assessed by comparing the actual and predicted grand mean spike counts, and only accepting units whose predicted grand mean spike counts fell in the range 0.83-1.2x of the actual values. Of 856 units, 661 units fulfilled this criterion.

Finally, we used the tuning parameters for each unit to synthesize the responses to 1000 images per condition. For each unit, we sampled x in Eq. 30 as 1000 draws from a uniform distribution between 0 and 1 and used those values to compute spike count rates, which were converted to spike counts by drawing from a Poisson process.

## Supplementary Information

**Fig. S1.**
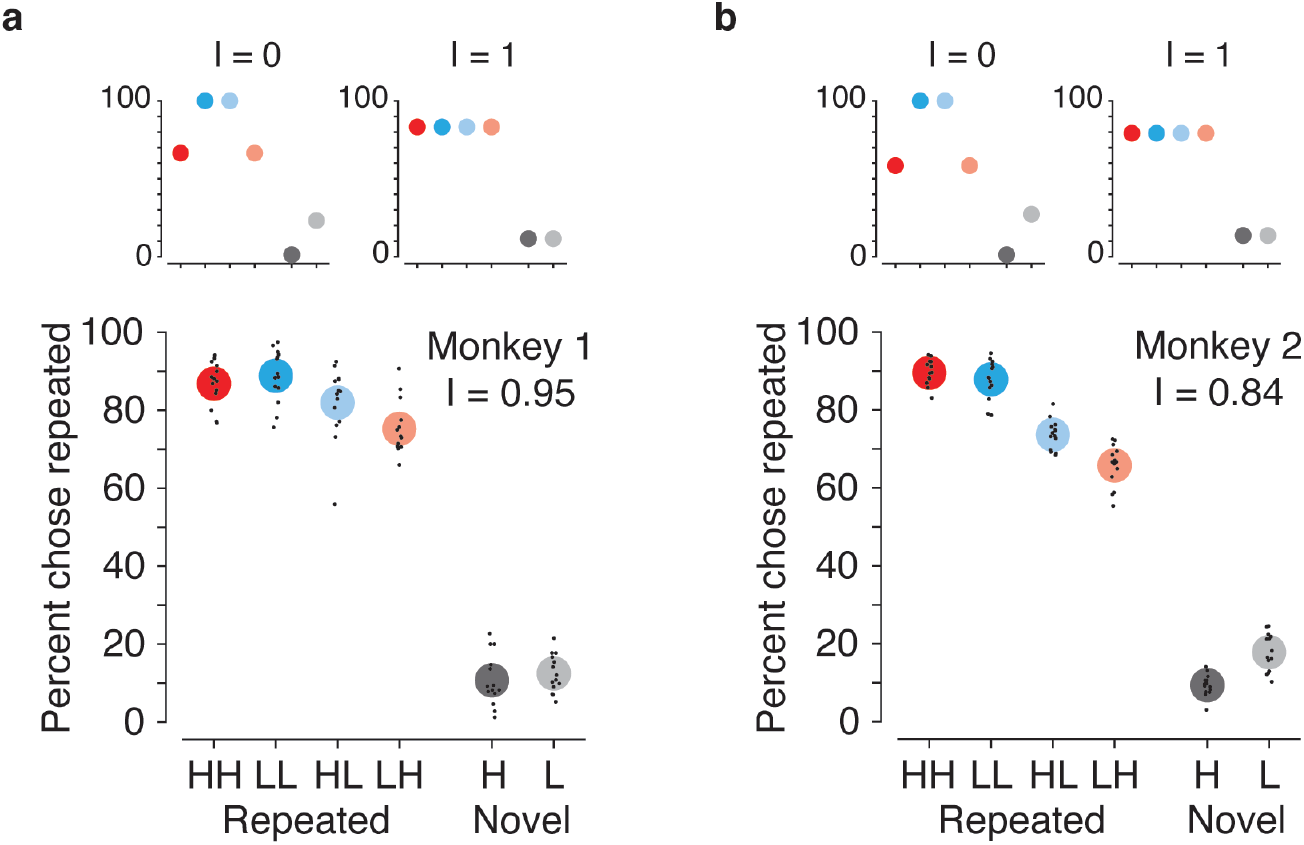
Behavioral performance patterns for individual monkeys. **(a-b)** Fig. 1c replotted for two animals. Small black dots indicate average performance for an individual session and large colored dots indicate the average performance across sessions (14 sessions per animal). The contrast invariance reflected in each behavioral pattern (I) is labeled in each plot. Insets correspond to behavioral patterns with maximal (I = 0) and minimal (I = 1) contrast confusion, matched for overall performance.

**Fig. S2.**
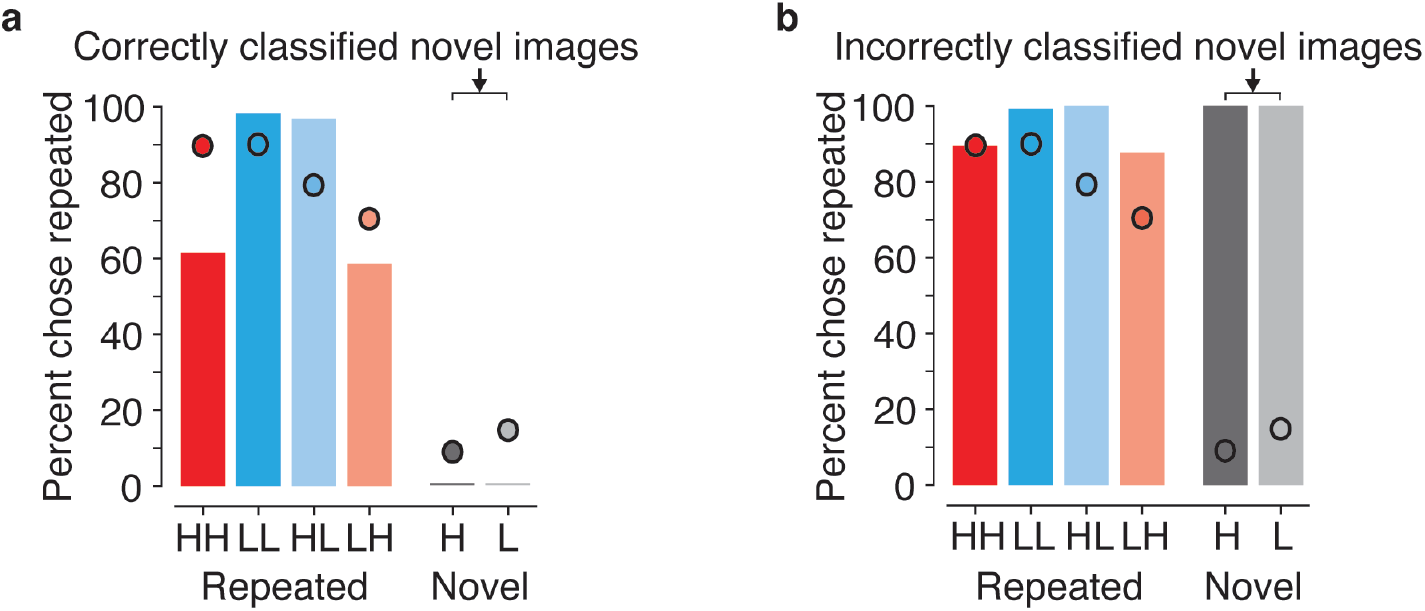
RS decoding performance, sorted by decoding performance on novel trials. Shown is a breakdown of the behavioral pattern predicted by the RS decoder depicted in Fig. 2a, sorted by images that were **(a)** correctly, and **(b)** incorrectly classified when presented as novel. These results indicate that even when novel images are correctly classified (panel a), the predicted behavioral pattern for repeated images reflects contrast confusions.

**Fig. S3.**
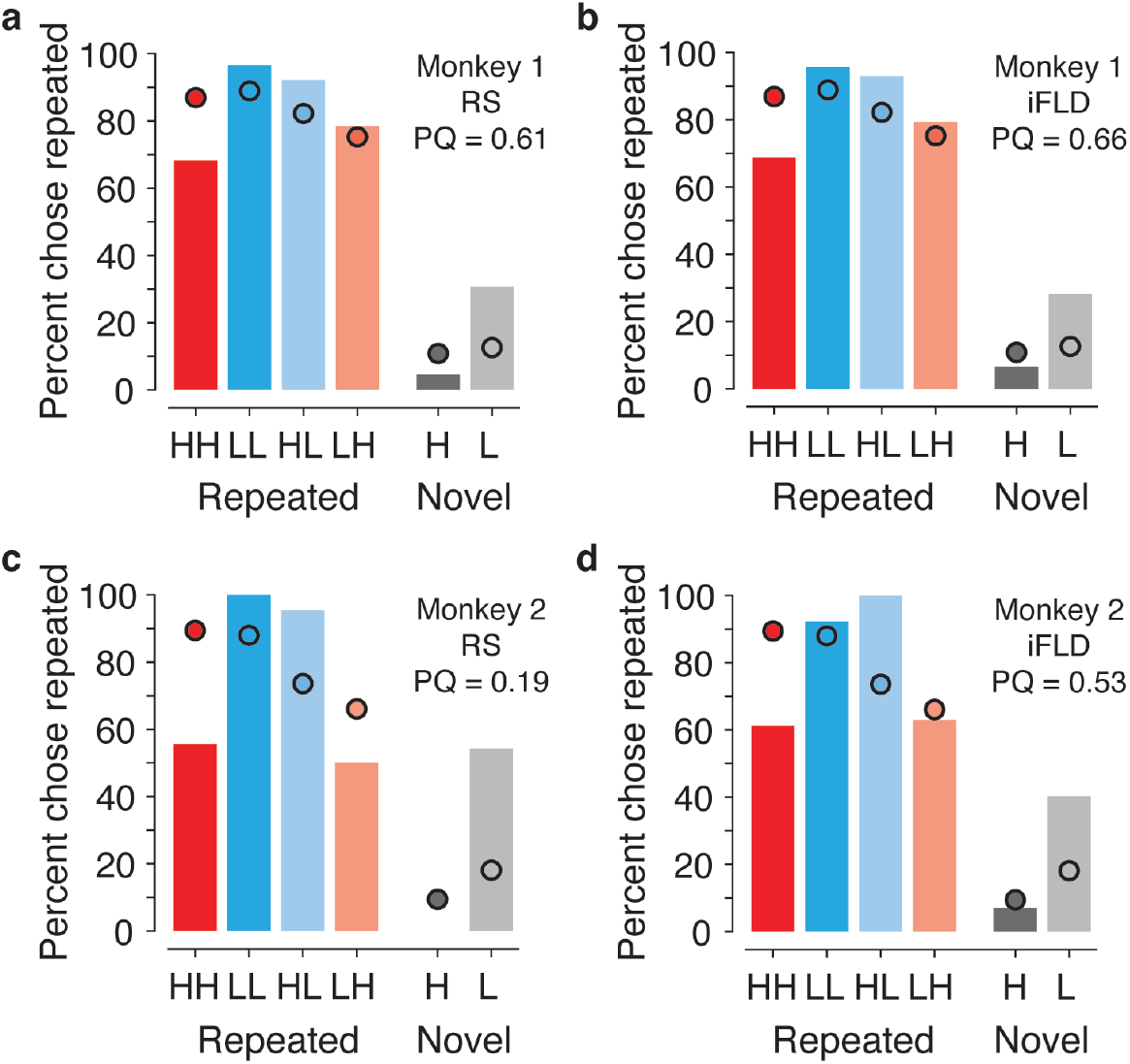
Classic linear decoders fail to map IT neural responses to behavior for each monkey. **(a, c)** Fig. 2a replotted for each animal. **(b, d)** Fig. 2b replotted for each animal. In all panels, dots indicate the actual behavioral patterns and bars indicate the neural predictions of behavior for each type of linear decoder. Prediction quality (PQ) is indicated for each case.

**Fig. S4.**
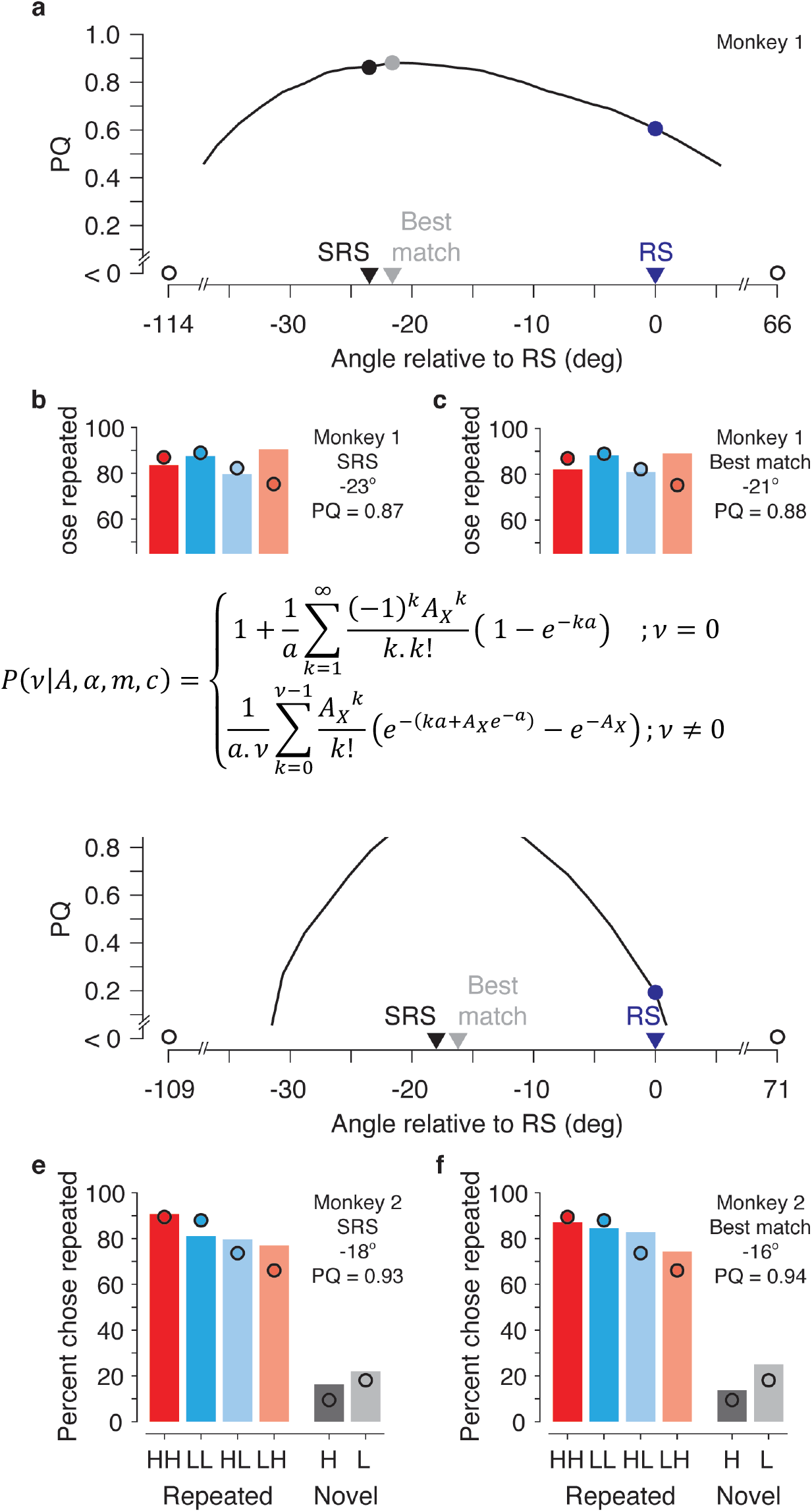
Neural predictions of behavior for a family of weighted linear decoders that include SRS, plotted for each monkey. **(a, d)** Fig. 3b, plotted for each animal: prediction quality (PQ) for the family of linear decoders that lie on the plane spanned by RS and contrast axis (Fig. 3a). Markers correspond to SRS (black), RS (blue), and the linear decoder with largest PQ (grey). **(b, e)** Fig. 3c, plotted for each animal: the alignment of the actual behavioral pattern and the SRS prediction **(c, f)** Fig. 3d, plotted for each animal: the alignment of the actual behavioral pattern and the decoder with the highest PQ on this plane. In b-f, dots indicate actual behavioral patterns and bars indicate the linearly decoded neural predictions of behavior. The decoder’s direction relative to RS, and prediction quality (PQ) are labeled for each case.

**Fig. S5.**
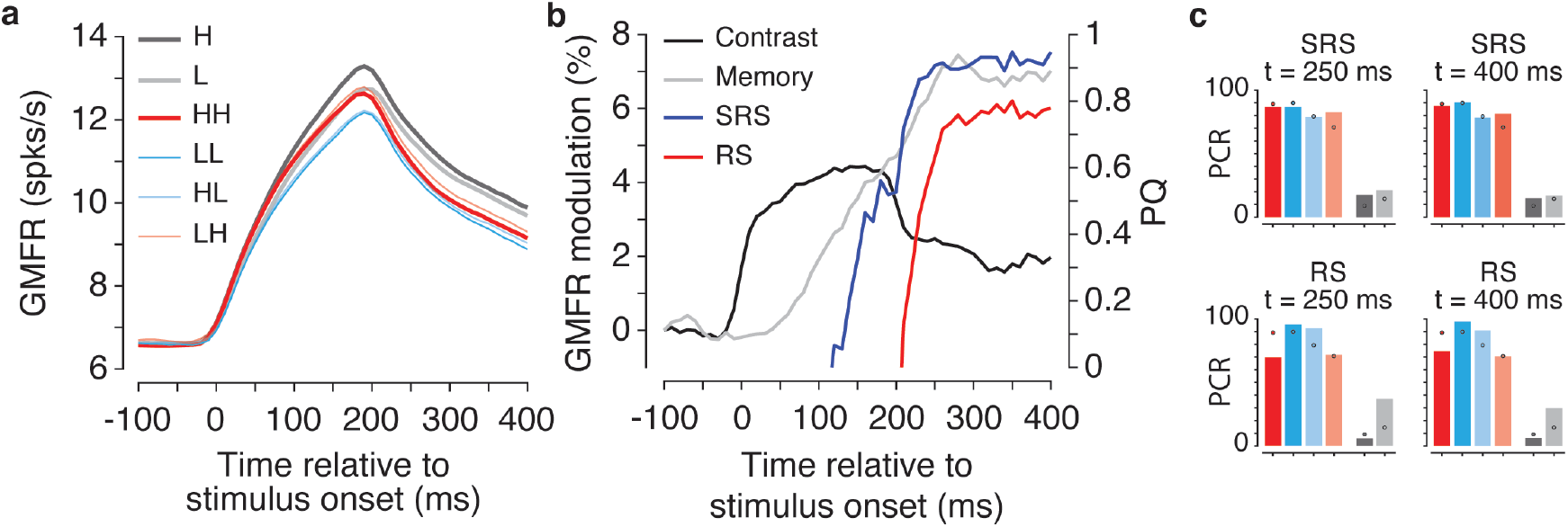
Temporal evolution of IT contrast and memory representations and their impact on decoding image memory. **(a)** Grand mean firing rate (GMFR) of IT neurons as a function of time relative to stimulus onset. Traces in different colors correspond to different conditions (n = 856 units). **(b)** The evolution of contrast modulation (black) and memory modulation (grey), plotted along the left y-axis. We used traces depicted by thick lines in (a) to compute the modulations, such that: 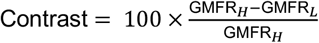, and 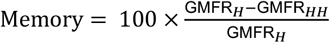, where GMFR denotes grand mean firing rate in the subscripted GMFR& condition. The time-course of prediction quality (PQ) for RS (red) and SRS (blue) decoders are also shown, plotted along the right y-axis. **(c)** The behavioral pattern predicted by RS (bottom row) and SRS (top row) decoders for two time points: t = 250 ms (first column), and t = 400 ms (second column). To perform these analyses, spikes were counted in 200 ms bins with bin centers that were shifted by 10ms.

**Fig. S6.**
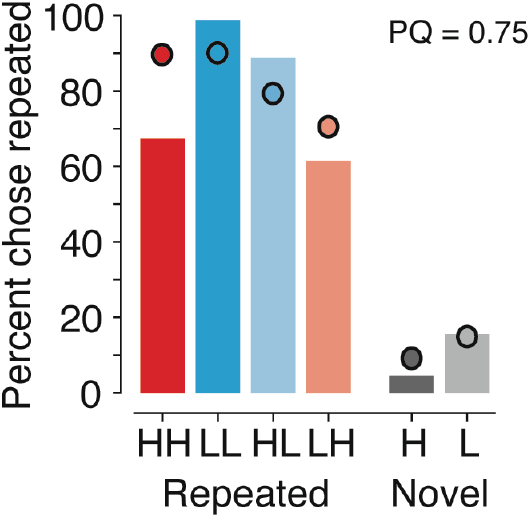
Variant decoder that incorporates a contrast correction. Shown are the results for a decoder that operates by correcting the projection of IT responses along the RS axis by subtracting an estimate of the mean population response across novel images in each contrast condition (see Methods). The estimate of the mean population response at each contrast was computed after classifying novel images by contrast based on the training data, using the same prototype linear decoder used for SRS (see Methods). This scheme produced a lower prediction quality (PQ = 0.75) than SRS (PQ = 0.88; Fig. 3c).

**Fig. S7.**
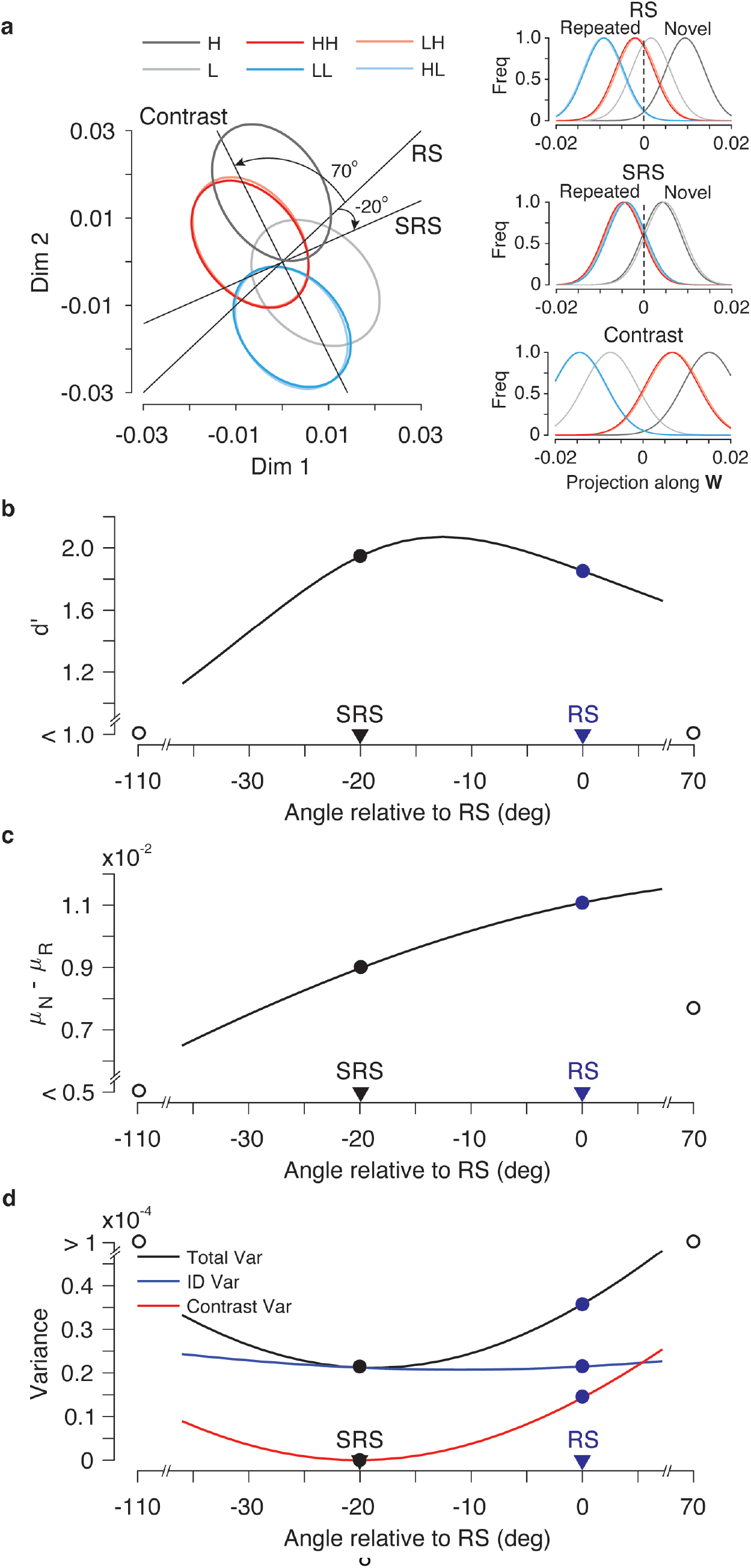
Synthetic data generated from the 4-parameter model recapitulates the actual data. Simulations were performed for 650 units x 4K images (1K images/condition). All analyses were performed in the same manner as described for the physiological data. Plotted for the synthetic data **(a)** Fig. 3a **(b-d)** Fig. 4b-d.

**Fig. S8.**
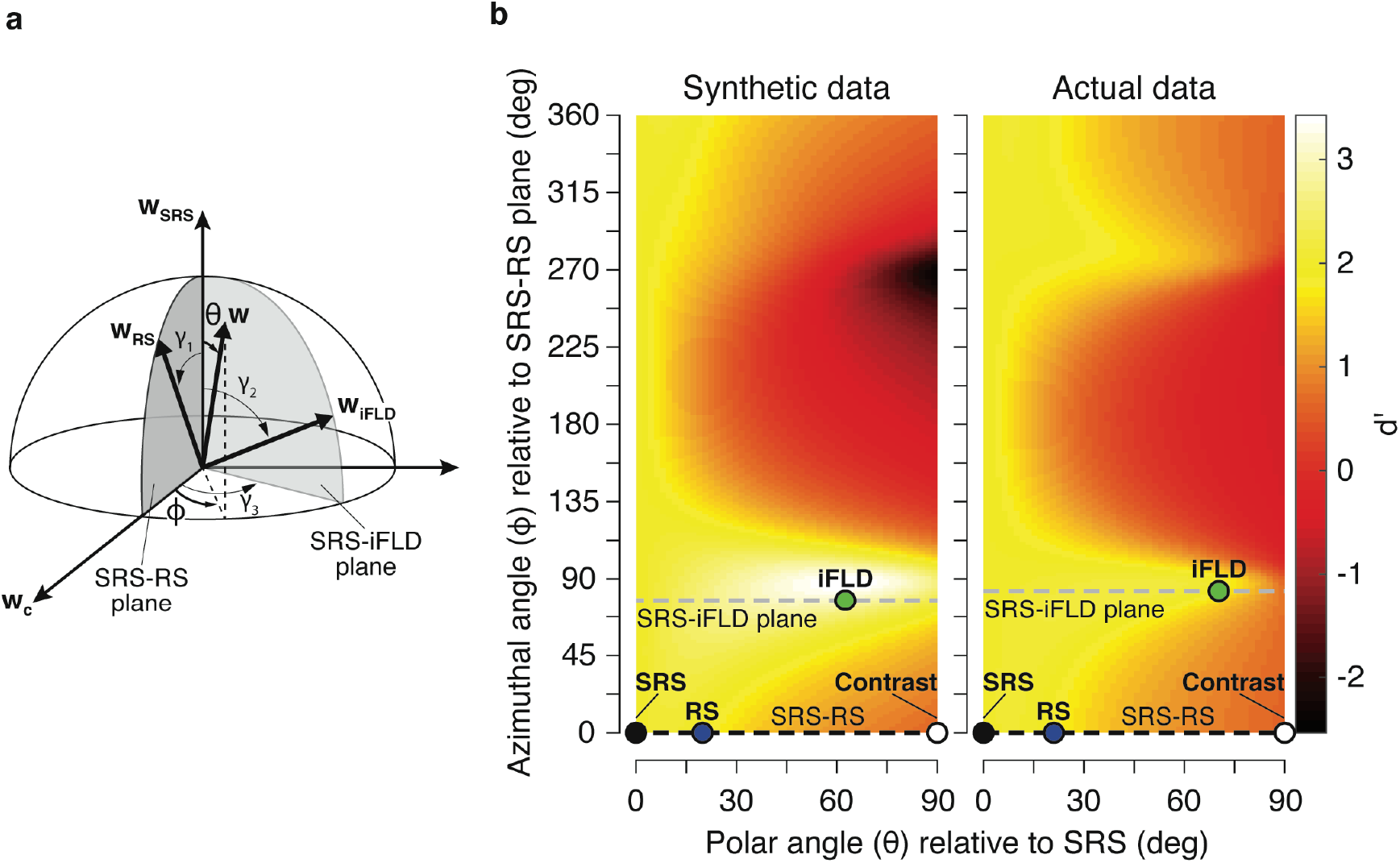
Decoding performance in the 3-D subspace spanned by the SRS, RS and iFLD linear decoders. **(a)** Depiction of the 3-D subspace in a spherical coordinate system. A decoding axis (**w**) in this subspace is determined by the polar angle relative to SRS (θ) and the azimuthal angle (ϕ) relative to the contrast axis in the 2-D plane defined by SRS and RS. Because the 3-D subspace spanned by SRS, RS, and iFLD is a non-orthogonal coordinate system, we used angles ***γ***_**1**_, ***γ***_**2**_, and ***γ***_**3**_ to transform the illustrated cartesian coordinate system to the coordinate system spanned by SRS, RS, and iFLD (see Methods). ***γ***_**1**_ indicates the angle between SRS and RS, ***γ***_**2**_ is the angle between SRS and iFLD, and ***γ***_**3**_ represents the angle between SRS-RS and SRS-iFLD planes (see Methods). **(b)** Discriminability for image memory (d’), computed as described for Figs. 4b, 5b, and *SI Appendix,* Fig. S7b, plotted as a function of polar and azimuthal angles in the 3-D subspace (see Methods). Shown are the results applied to synthetic data taken from the model described for Fig. 5 and the actual data. Values corresponding to SRS, RS, iFLD, and contrast are labeled by black, blue, green, and open markers, respectively. Black and grey dashed lines mark the azimuthal angles associated with SRS-RS and SRS-iFLD planes, respectively.

